# Mutational scanning pinpoints distinct binding sites of key ATGL regulators in lipolysis

**DOI:** 10.1101/2023.05.10.540188

**Authors:** Johanna M. Kohlmayr, Gernot F. Grabner, Anna Nusser, Anna Höll, Bettina Halwachs, Evelyne Jany-Luig, Hanna Engelke, Robert Zimmermann, Ulrich Stelzl

**Affiliations:** Institute of Pharmaceutical Sciences, Pharmaceutical Chemistry, University of Graz, Austria; Institute of Molecular Biosciences, Biochemistry, University of Graz, Austria; Field of Excellence BioHealth - University of Graz, Austria; BioTechMed-Graz, Austria

**Keywords:** protein interaction perturbation, deep scanning mutagenesis, MAVE, yeast two-hybrid, lipid metabolism

## Abstract

ATGL is the key enzyme in intracellular lipolysis playing a critical role in metabolic and cardiovascular diseases. ATGL is tightly regulated through a known set of protein-protein interaction partners with activating or inhibiting functions in control of lipolysis. However, the binding mode and protein interaction sites of ATGL and its partners are unknown. Using deep mutational protein interaction perturbation scanning we generated comprehensive profiles of single amino acid variants effecting the interactions of ATGL with its regulatory partners: CGI-58, G0S2, PLIN1, PLIN5 and CIDEC. Twenty-three ATGL variants gave a specific interaction perturbation pattern when validated in co-immunoprecipitation experiments in mammalian cells. We identified and characterized eleven, highly selective ATGL “switch” mutations which affect the interaction of one of the five partners without affecting the others. Switch mutations thus provided distinct interaction determinants for ATGL’s key regulatory proteins at an amino acid resolution. When tested for triglyceride hydrolase activity *in vitro* and lipolysis in cells, the activity patterns of the ATGL switch variants traced to their protein interaction profile. In the context of structural data, the integration of variant binding and activity profiles provided important insights into lipolysis regulation and the impact of mutations in human disease.

## Introduction

Lipid droplets (LDs) are the main cellular storage organelles for triacylglycerol (TAG). The dysregulation of the TAG metabolism is associated with human metabolic pathologies, including obesity, cardiac pathophysiology, non-alcoholic fatty liver disease, lipodystrophy and neutral lipid storage disease (NLSD) (Olzmann and Carvalho, 2019; Grabner et al., 2021). The highly conserved process of intracellular lipolysis mediates the hydrolysis of TAG stored in LDs. Adipose triglyceride lipase (ATGL; gene name: *PNPLA2*) is the key enzyme in lipolysis and catalyzes the first step of TAG breakdown by cleaving off the first fatty acid from the TAGs resulting in a diacylglycerol (DAG) and free fatty acids (Zimmermann et al., 2004). The catalytic activity of ATGL is important for the regulation of whole-body energy homeostasis and lack of functional ATGL leads to severe metabolic phenotypes. Global ATGL-deficient mice accumulate TAG in most tissues, exhibit reduced plasma fatty acid concentrations, improved glucose tolerance, and insulin sensitivity, but die prematurely due to cardiomyopathy (Haemmerle et al., 2006). Rescue studies, expressing ATGL exclusively in the heart and thus preventing cardiomyopathy (Schoiswohl et al., 2010; Haemmerle et al., 2011), protected mice from high fat diet (HFD)-induced obesity (Schreiber et al., 2015; Schweiger et al., 2017). Adipocyte-specific deletion of ATGL in mice (Schoiswohl et al., 2015) had similar molecular and phenotypic effects as observed in the rescue studies, implying that ATGL inhibition in adipose tissue leads to a beneficial metabolic phenotype. Similarly, pharmacological inhibition of ATGL with small molecule inhibitors represents a viable treatment strategy for obesity-associated disorders and cardiac insufficiency (Schweiger et al., 2017; Grabner et al., 2022).

Activation of the lipolysis pathway in adipocytes occurs through β-adrenergic GPCR signaling for example by catecholamines. Lipolysis, including ATGL activity, is post-translationally regulated by proteins located at the LD surface (Cerk et al., 2018). The current model (Grabner et al., 2021) proposes that under basal conditions members of the perilipin family coat the LD surface thereby restricting the access of ATGL to the LDs. At the same time, Perilipin 1 (PLIN1) sequesters the activating protein CGI-58 (gene name: *ABHD5*) preventing its association with and activation of ATGL. Upon stimulation, PKA-dependent phosphorylation of PLIN1 and CGI-58 leads to dissociation of the complex and CGI-58 release, which then binds and activates ATGL (Granneman et al., 2009; Sahu-Osen et al., 2015). Additionally, PKA phosphorylation also recruits hormone sensitive lipase (HSL), the second lipase required for DAG cleavage in the lipolysis pathway, from the cytosol to LDs (Recazens et al., 2021). However, this model is incomplete as studies showed that TG hydrolysis does not require PKA-dependent activation of HSL (Rondini et al., 2017) and that a S239D phospho-mimicry variant of CGI-58 does not require PLIN1 phosphorylation for β-adrenergic stimulated ATGL activation (Sahu-Osen et al., 2015). Together, published data suggest that additional phosphorylation targets are elusive and that the order of molecular events in control of the lipolysis pathway is not fully understood. Detailed knowledge of protein-protein interaction (PPI) determinants is prerequisite for a full mechanistic understanding of ATGL regulation and ATGL-dependent lipolysis mechanism.

One of the most important ATGL PPI partners (**Figure 1A**) is CGI-58, the co-factor required for full hydrolytic activity of ATGL (Lass et al., 2006). Previously, structural protein domains required for the PPI have been identified (Schweiger et al., 2008; Cornaciu et al., 2011; Tseng et al., 2022), but specific ATGL amino acid residues or binding surfaces remained undetermined. Conversely, the interaction of G0S2 (G0/G1 switch gene 2) with ATGL potently inhibits ATGL TAG hydrolase activity (Yang et al., 2010). Mutational analysis of a G0S2 peptide comprising amino acid residues 20 to 44 defined the minimal G0S2 binding site for the interaction with the N-terminal patatin domain of ATGL (Yang et al., 2010) (**Figure 1A**). However, introducing single amino acid mutants in the full-length G0S2 protein did not disrupt the PPI in co-IP experiments and only slightly altered the inhibitory effect of G0S2 on ATGL activity, indicating that additional parts of G0S2 participate in this interaction (Riegler-Berket et al., 2022). Interaction determinants on the side of ATGL protein as well as the inhibitory mechanism are unknown. CIDEC (cell death inducing DFFA like effector c, also FSP27), exclusively found in white and brown adipose tissue, is another inhibitory protein of ATGL (Puri et al., 2007). Interaction between ATGL and CIDEC inhibits ATGL-mediated lipolysis but does not directly affect ATGL’s TAG catalytic activity (Yang et al., 2013; Grahn et al., 2014). The interaction is mediated via amino acid residues S120 to P220 of CIDEC (Grahn et al., 2014) (**Figure 1A**). Perilipins, such as PLIN1 and PLIN5, are major inhibitory regulators of lipolysis under basal conditions (Najt et al., 2022) by binding of CGI-58 (Subramanian et al., 2004; Yamaguchi et al., 2004; Granneman et al., 2009) and by restricting the access of lipases to the LD surface. Plin1 deficiency leads to increased lipolysis, hence Plin1 knock-out mice are lean, show reduced LD size, are resistant to HFD-induced obesity, and develop insulin resistance (Tansey et al., 2001). Interestingly, also upon overexpression of Plin1, adipose tissue is reduced and mice become resistant to HFD-induced obesity (Miyoshi et al., 2010; Sawada et al., 2010). Like Plin1, Plin5 promotes TAG accumulation under basal condition but increases lipolysis in response to PKA stimulation (Wang et al., 2011; Pollak et al., 2015). Plin5 directly binds ATGL via residues R417 to F463 (**Figure 1A**) (Granneman et al., 2011; Wang et al., 2011). Through its very C-terminus, Plin5 also directly interacts with mitochondria linking LDs to FA oxidation (Wang et al., 2011; Kien et al., 2022). Heart-specific knockout of Plin5 in mice partially phenocopies ATGL loss of function (Kuramoto et al., 2012; Pollak et al., 2013).

**Figure 1:**
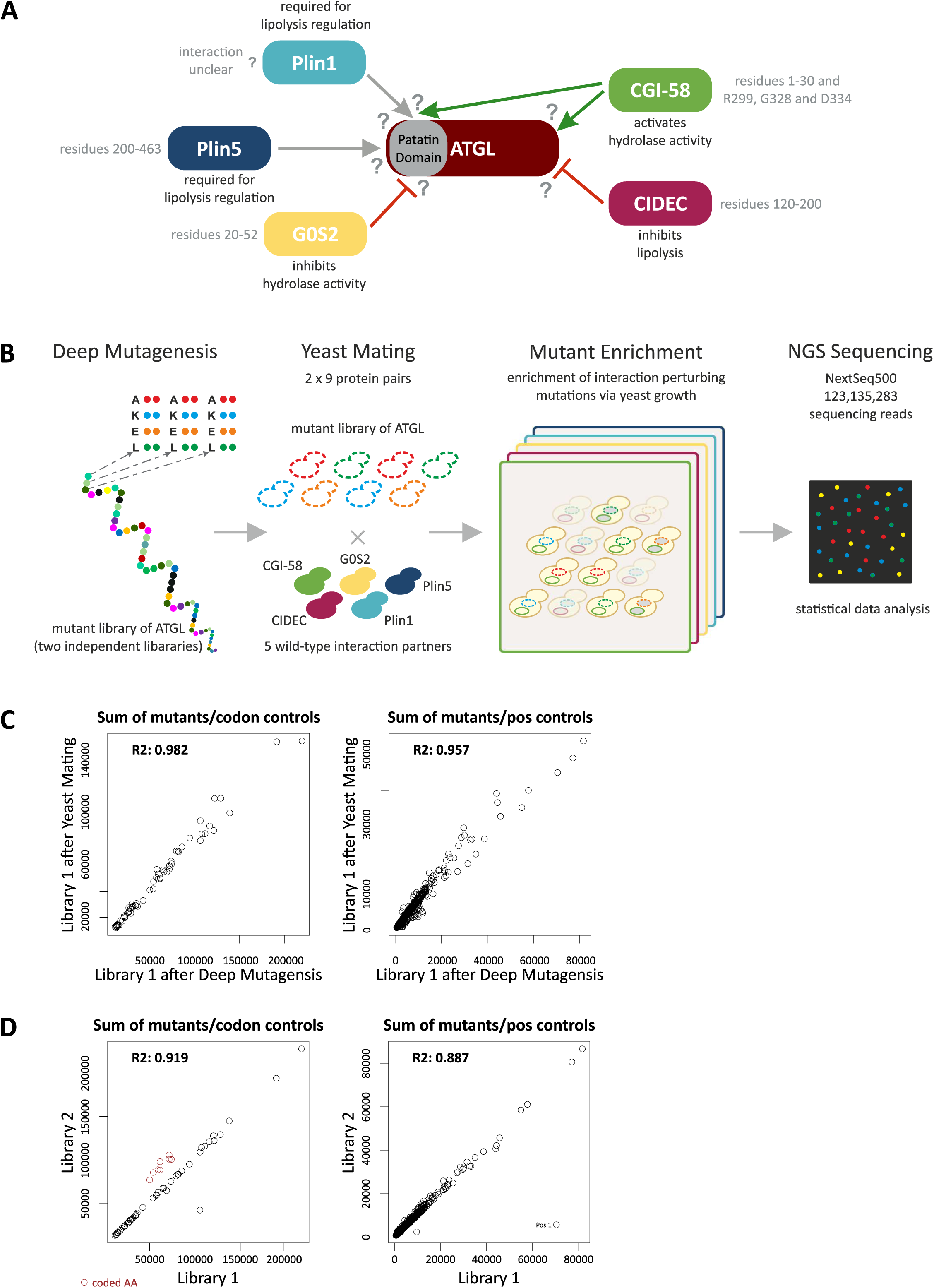
Mapping ATGL protein interaction determinants using deep mutational interaction perturbation scanning. **A.** Schematic overview of ATGL and its interaction partners. Known AA residues mediating the interactions are annotated, see text for references. The Perilipin family members bind ATGL within the patatin domain. G0S2 also binds ATGL within the patatin domain, inhibits its hydrolytic activity. CIDEC, another lipolysis inhibiting protein interacts with the C-terminal part of ATGL. CGI-58 binds ATGL via the patatin domain as well as the C-terminal part. However, no interactions sites on ATGL for its partners were identified. **B**. Schematic overview of the deep mutational interaction perturbation screening approach. Plasmid libraries of ATGL were generated using array programmed deep mutagenesis, exchanging single amino acid to A, K, E or L, respectively. Reverse Y2H strains were transformed with the ATGL libraries and mated with yeast strains expressing a WT interaction partner. Interaction disrupting mutations were enriched through yeast growth, and mutations perturbing the PPI were identified by NGS sequencing. **C.** Scatter plot of read counts of library 1 in the Gateway Entry vector after deep mutagenesis with the library after yeast mating prior to growth selection. Read counts compared for sum of mutants per codon (left) as well as per position (right). R2 values indicate that the library did not change during the cloning procedure. **D.** Scatter plot of read counts of the two independent ATGL libraries. Library 1 and library 2 were compared for their sum of mutants observed per codon (left) as well as per position (right). Red colored circles indicate programmed AKEL mutations which are higher in library 2 than library 1. The outlier data point (Position 1) in library 1 in the right graph likely stems from an PCR amplification step early during preparation.

Deep mutational scanning approaches that assess the functional effects of thousands of variants of a protein have become a prime tool in functional genomics (Shendure and Fields, 2016). They allow to study the impact of every possible single amino acid substitution in a protein on its function, protein stability, biophysical properties, and protein-protein interactions as well as a functional annotation of disease-associated human variants (Kunowska and Stelzl, 2022). Yeast based protein-protein interaction assays, such as yeast protein fragment complementation or yeast two-hybrid assays, that enable the enrichment of interacting protein variants or non-interacting variants, respectively, are key to deep mutational protein interaction scanning (Moesslacher et al., 2021). The assays have been used to scan several key protein interactions, including PPIs of the BBSome complex (Woodsmith et al., 2017), the interaction between BARD and BRCA1 (Starita et al., 2015), leucine zipper PPIs (Diss and Lehner, 2018), the Ras-Raf pair (Hidalgo et al., 2022; Weng et al., 2022), peptide binding of a SH3 and a PDZ domain (Faure et al., 2022) and a set of NF2 tumor suppressor protein interactions (Moesslacher et al., 2022). We apply reverse yeast two-hybrid (rY2H) growth selection for efficient enrichment of non-interacting mutant protein versions from comprehensive single amino acid protein variant libraries (Woodsmith et al., 2017). A quantitative sequencing readout identifies amino acid substitutions that negatively impact the capacity to interact with a protein partner (**Figure 1B**). When assaying a deep mutational pool of a protein with multiple protein interaction partners, we can infer mutations that selectively impact the binding of individual partners rather than mutations that impair the function of the protein as a whole.

Here, we use the deep mutational interaction perturbation approach to comprehensively chart the impact of >4000 single amino acid substitutions in ATGL on the binding to its five most important interaction partners, CGI-58, G0S2, Plin1, Plin5 and CIDEC. The results of our deep mutational interaction scanning experiments first of all define contact sites for the interacting proteins that govern ATGL activity and function. Moreover, we specifically pinpoint a set of eleven switch mutations that selectively impair the interaction of ATGL with one of the five protein partners only. These switch mutations show defined functional patterns *in vitro* and lipolysis phenotypes in cells. Definition of amino acid resolution binding determinants for ATGL regulatory proteins provides mechanistic insights and novel hypotheses on the mechanism of regulated intracellular lipolysis.

## Results and Discussion

### Deep mutational scanning interaction perturbation of ATGL-PPIs

We first established the interactions between full length human ATGL and CGI-58, G0S2, PLIN1, PLIN5 and CIDEC in our Y2H assay system (Worseck et al., 2012; Weimann et al., 2013) (**Figure 1A**). Applying the nicking mutagenesis protocol of Wrenbeck *et al*. (Wrenbeck et al., 2016), two libraries of ATGL variants were constructed. These two libraries contained mutations of each and every amino acid against alanine (A::GCT and A::GCA), lysine (K::AAA and K::AAG), glutamic acid (E::GAA and E::GAG) and leucine (L::TTG and L::CTA), using two different codons per amino acid exchange (**Figure 1B**). This totaled up to 4000 single amino acid mutations in each ATGL cDNA library. During the subcloning into Y2H vectors, the colony number of the libraries was kept above 1 million, exceeding the theoretical number needed to maintain each ATGL variant in the library by 2 orders of magnitude. Sequencing of the two libraries confirmed that the number of mutations per codon and mutations per amino acid positions in ATGL did not change during the cloning procedure (library after mutagenesis vs after yeast mating, **Figure 1C**). Read counts of non-programmed and programmed AKEL mutations between the two libraries were highly correlated with library 2 having a slightly higher rate of programmed mutations than library 1. Except amino acid position 1 no positional bias was observed (**Figure 1D**).

The two ATGL variant libraries were each tested with the five wild-type interaction partner proteins in the perturbation screen (**Figure 1B**). Additionally, wild-type protein ORFs were used in different Y2H vector configuration (**Suppl. Table 1**) so that two (PLIN1, CIDEC), four (CGI-58, PLIN5) and six (G0S2) screens were combined into one ATGL perturbation profile for each interaction partner (**Figure 2A**). After sequencing, relatively low cutoffs were applied to generate the interaction profiles, which is reflected by many mutation hits distributed in the condensed profiles for the five proteins (**Suppl. Table 2** and **Suppl. Table 3**). Across the whole data set, mutations of proline, glycine, cysteine as well as threonine and serine contributed most frequently to the interaction disrupting mutations, which may reflect the importance of those amino acids for folding and stability (Diss and Lehner, 2018; Faure et al., 2022) (**Figure 2B**). We detected many variants that impacted several protein interactions (full data set as **Suppl. Table 3**), however, mutations in the region around amino acid 250, a linker region between the patatin domain and the C-terminal part of ATGL, hardly had any impact on binding of the interaction partners. In a close inspection of the profiles, we focused on identifying positions where mutations selectively impaired binding of one or two proteins, while having little effect on the other interaction partners. For example, N39E had a high score for CGI-58 and G0S2 but not for the PLINs or CIDEC (**Figure 2C**). P103K was enriched in the screen with PLIN1 and PLIN5, but not with the other three interaction partners. The mutational profile for L159 showed that, besides a score for CGI-58 and G0S2, L159A most strongly reduced binding to PLIN5. Evaluation of the variant profiles led to the prioritization of 52 ATGL variants for further individual assessment of binding behavior (**Suppl. Table 4**).

**Figure 2:**
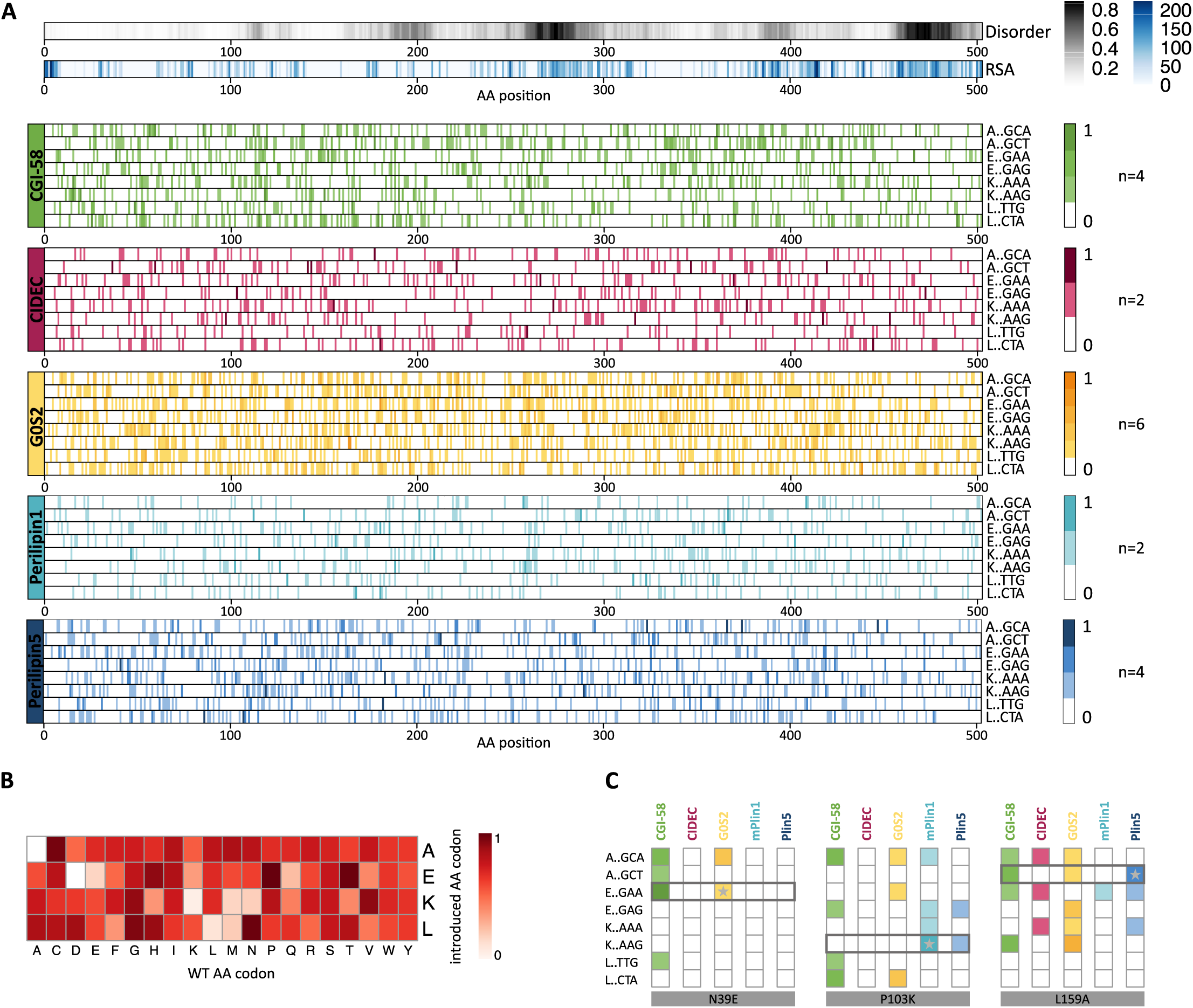
Interaction perturbation maps of ATGL with five key regulatory binding partners. **A.** Combined interaction perturbation profiles for each ATGL binding partner. Top two profiles indicate relative solvent accessibility (RSA) and disorder obtained from IUPRED over the ATGL sequence. Combined, normalized individual profiles at codon resolution for the programmed AEKL mutations for the five interaction partners are shown (n number of experiments). See also Suppl. Table 3. **B.** Heat map of frequency of single amino acid substitutions across all interactions. AKEL substitutions were counted for every amino acid across all probed interactions and normalized to the frequency of occurrence in the ATGL wild-type sequence. **C.** Zoom in at the perturbation profiles showing binding specificities of amino acid residues N39, P103 and L159. Mutations at position N39 affected the interaction of ATGL with CGI-58 and G0S2. The variant N39E (E::GAA) was selected as for further validation. Mutations at P103 influence several interactions with ATGL, however the exchange to K (K::AAG) selectively perturbed the interactions with Plin5 and Plin1. The L159A (A::GCT) substitution strongly perturbed the interaction with Plin5 and was selected for detailed variant analysis.

### Co-immunoprecipitation experiments define ATGL switch variants

Next, we tested the specificity of the mutational effects on individual protein interaction partners using a luminescence-based mammalian co-immunoprecipitation assay in HEK293T cells (LUMIER) (Hegele et al., 2012; Taipale et al., 2013). Protein A was fused to the N-terminus of the ATGL variant protein set, comprising 52 single amino acid ATGL mutants as well as ATGL wild-type and a non-binding control protein. The Proteins were co-expressed with wild-type CGI-58, G0S2, CIDEC, Plin1 and Plin5, as firefly-luciferase fusion proteins. The HEK293T cell lysates with the transiently expressed protein pairs were subject to immuno-precipitation in immunoglobulin-G coated microtiter plates and co-immunoprecipitation of the interaction partners was quantified as luminescence signal. Hence, we comprehensively tested all five interaction partners pairwise with 54 ATGL constructs in ∼ 260 co-immunoprecipitation experiments for interaction in mammalian cells. To exclude that binding was substantially influenced by protein expression in the mammalian cell lines, similar expression levels of ATGL mutants was confirmed by western blot analysis. Normalized to wild-type ATGL binding, log2 fold change binding values were calculated from two independent experiments, performed in triplicates (**Figure 3A, Suppl. Table 5**). Compared to wild-type ATGL, 29 variants did not show statistically significant differences in binding to any of the five interaction partners (**Figure 3B**). This result is consistent with different assay sensitivities between the deep scanning sequencing redout and the co-immunoprecipitation experiments. Conversely, the G189E mutant, while expressed as soluble protein of the correct size, perturbed the interaction with all interaction partners, an effect on binding considered non-informative. Importantly, 22 ATGL variants impacted individual protein interactions in a specific manner.

**Figure 3:**
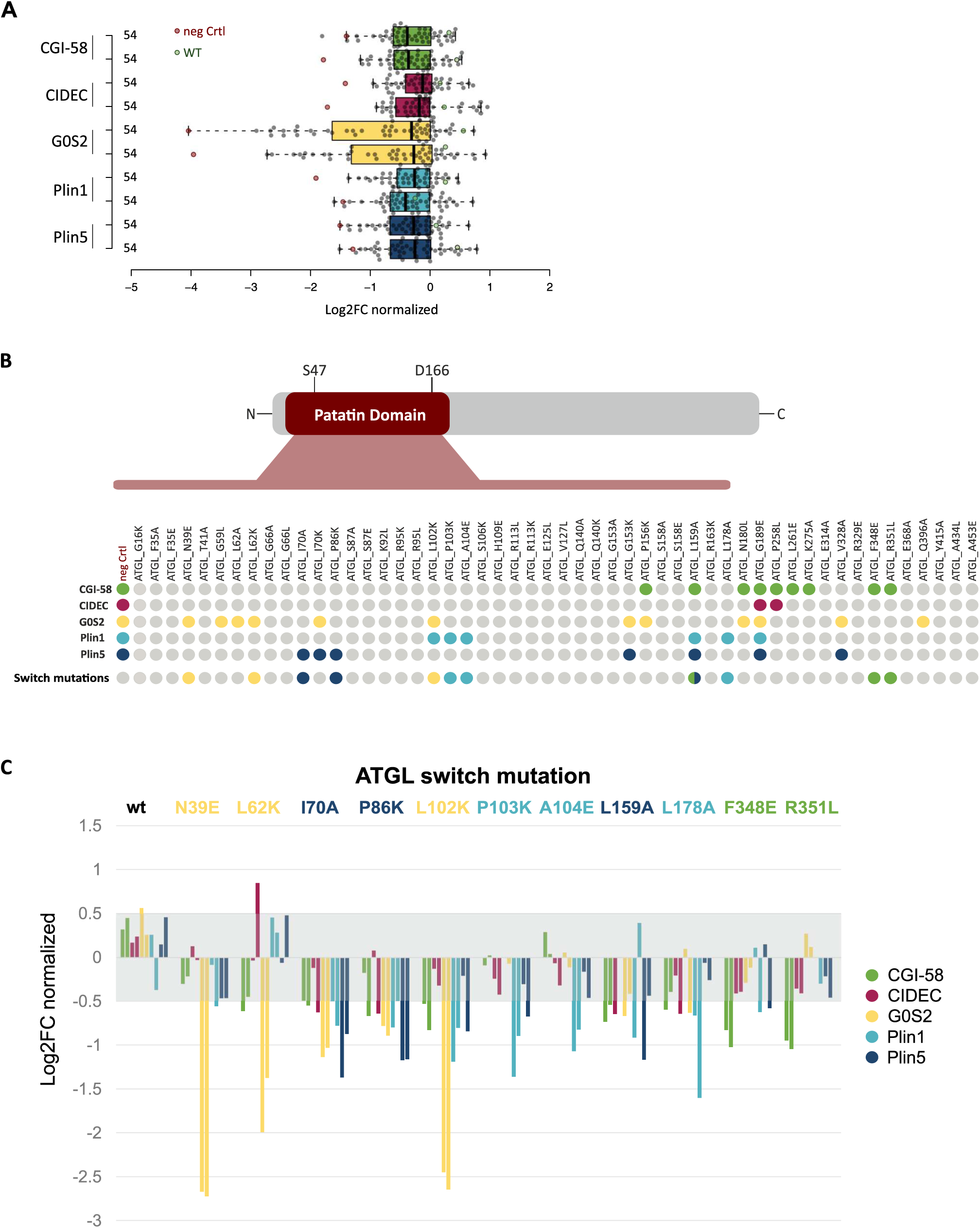
ATGL switch mutations selectively interrupt individual binding partners. **A.** Luciferase based co-immunoprecipitation of 52 ATGL mutants with five protein interaction partners. Box plot of showing the mean of triplicate measurements of two experiments each probing murine Plin5 (blue), murine Plin1 (cyan), G0S2 (yellow), CIDEC (red) and CGI-58 (green) for interaction with the human ATGL variants (gray data points), ATGL wild-type (green data points) and a negative control (red data points). Log2 fold changes in binding relative to ATGL wild-type was determined within every experiment (Suppl. Table 5). Values were normalized across the experiments to the 3^rd^ quartile indicating increased or decreased interaction signal. **B.** Systematic overview of the binding data of the 52 ATGL variants. Variants are shown with their location within the patatin domain or in the C-terminal half of the protein. ATGL variants reducing the binding by more than twofold are marked as colored dots according to the affected interaction partner. The switch mutants are highlighted separately. Color code: CGI-58 green, CIDEC red, G0S2 yellow, mPlin1 cyan and mPlin5 blue. **C.** Binding data for eleven ATGL switch mutations. For each interaction partner (color coded) normalized log2FC of two experiments performed in triplicates are shown. Variants are color coded according to the most specific binding alteration with a single partner, log2FC values within −0.5 – 0.5 (gray area) indicate wild-type like binding behavior.

By virtue of its high binding signal, G0S2 was affected most strongly and by the highest number of ATGL variants, 12 in total. CIDEC on the other hand bound robustly to ATGL with only one mutation apart from G189E, namely P258L in the C-terminal part of ATGL, resulting in significantly decreased binding. This is in agreement with literature data that suggest the CIDEC binds to the C-terminal part of ATGL (Grahn et al., 2014). All other tested proteins had at least one interaction site in the patatin domain (**Figure 1A** and **3B**). As the majority of ATGL variants did not affect CIDEC binding, it may be regarded as positive control for the variants when tested with other binding partners in this comprehensive assay. For CGI-58 we validated nine variants that decreased ATGL interactions. Interestingly, five of the mutations were located in the C-terminal part of the ATGL protein outside the patatin domain. We also validated 11 variants that impacted Plin1 and/or Plin5 interaction with ATGL. In particular, we considered ATGL mutations that disrupted the interaction with one partner only, leaving the binding of all other unaffected. Such ATGL mutations were termed interaction switch mutations and they will likely have the most specific perturbation effect on ATGL regulation and signaling. We defined a set of 11 ATGL switch variants from our extensive binding data, three each for G0S2, PLIN1 and PLIN5 and two for CGI-58 (**Figure 3C**). Except for the two CGI-58 specific F348E and R351L mutations, all switch mutations were located within the patatin domain of ATGL that also contains the catalytic lipase center (**Figure 3B** and **C**).

The ATGL variants N39E, L62K and L102K showed a strong decrease in binding of the inhibitory protein G0S2, while other interaction partners were hardly affected. The L102K variant also decreased Plin1 binding, however the impact on G0S2 was much more substantial. Notably, Plin1 specific switch mutations P103K, A104E and L178A, did not impact Plin5 binding. The strongest reduction in binding of Plin5 was observed with the I70A, P86K and L159A ATGL variants (**Figure 3C**). The latter variant also affected binding of CGI-58. The interaction with CGI-58 was disrupted by nine mutations in total, however, F348E and R351L were identified as two highly specific CGI-58 switch mutations in the C-terminal part of ATGL (**Figure 3C**). In summary, we defined two to three ATGL switch mutations for its canonical regulatory interaction partners, CGI-58, G0S2, Plin1 and Plin5.

### ATGL switch variant binding correlates with TAG hydrolase activity profiles

The identification of ATGL interaction switch mutations, which are highly selective for one interaction partner, implied that the protein variant is at least partially functional. Still, it cannot be excluded that the switch mutations have an impact on protein function, such as lipid hydrolase activity. To address this, TAG hydrolase (TGH) activity of the 11 single amino acid switch mutant proteins was assessed *in vitro* (**Figure 4**).

**Figure 4:**
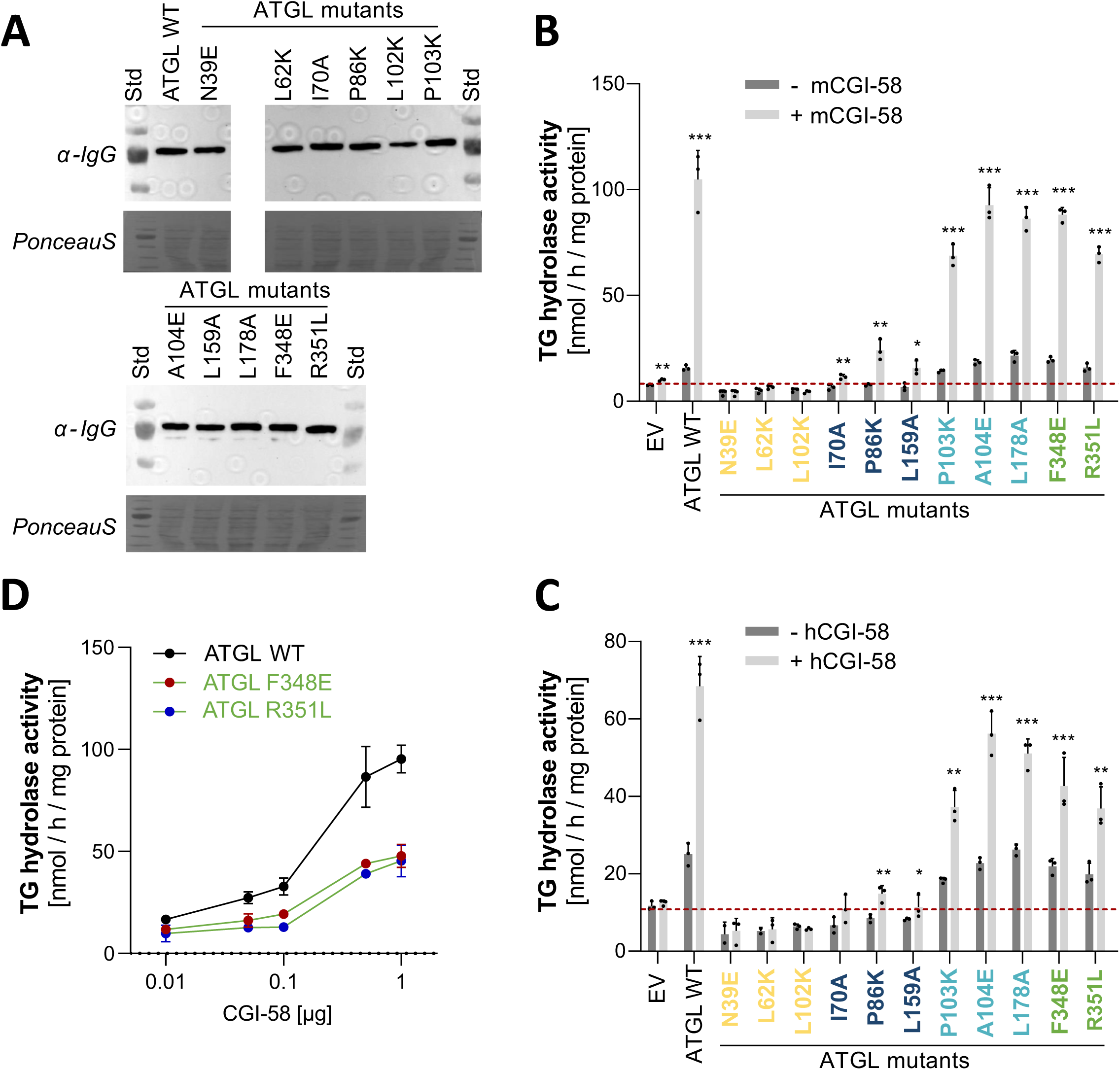
Enzymatic activity of ATGL switch variants. **A.** Western blot analysis of ATGL switch variant expression in Expi293 cells. 35 µg protein of the whole cell lysate was separated on a 10% SDS-gel and stained with anti-IgG-Ab. Bottom: Ponceau S staining of the blot membrane as loading control. **B-C.** TAG hydrolase activity detected in lysates of Expi293 cells. Activity was determined in the absence (-CGI-58, dark gray bars) or presence (+ CGI-58, light gray bars) of purified full-length mouse CGI-58 from E. coli (**B**) or 10µg lysate from cells expressing full-length human CGI-58 (**C**). EV, lysate with empty vector control, for measures of basal activity (-CGI-58, dark gray bars), the CGI-58 preparation was heat inactivated before addition. All samples were measured in triplicates and are shown as mean + standard deviation. Statistical significance was determined by unpaired two-tailed t-test (*p<0.005, **p<0.01, ***p<0.001). **D.** Dose dependent CGI-58 stimulation of ATGL switch variants. Data are presented as means, error bars indicate +/- standard deviation of triplicate experiments.

Wild-type ATGL and mutant ATGL versions were expressed in Expi293 cells, a mammalian system optimized for protein expression. ATGL protein expression was monitored via western blot analysis showing comparably high expression levels of all variants in the soluble lysate (**Figure 4A**). TGH activity of ATGL variants was determined in cell lysates with a PC/PI-emulsified radioactive isotope-labeled triolein substrate (Schweiger et al., 2014), and lysates of empty vector (EV)-transfected cells were used as negative control. Wild-type and all ATGL switch mutants were assayed for basal and CGI-58 stimulated hydrolase activity. Wild-type ATGL showed basal hydrolase activity that increased several-fold by stimulation with purified mouse CGI-58 protein (**Figure 4B**) and cell lysates containing full length human CGI-58 (**Figure 4C**). Stimulation with either purified CGI-58 or cell lysates containing human CGI-58 resulted in the same activity patterns across all variants.

The TGH activities of mutant ATGL proteins can be grouped into, *i)* inactive mutants, *ii)* mutants with reduced hydrolase activity, *iii)* mutants that had the comparable activity as wild-type ATGL, and *iv)* mutants with reduced response to CGI-58 stimulation. Inactive mutants N39E, L62K and L102K were insensitive to CGI-58 stimulation and the basal activities were indistinguishable from the empty vector control. The mutants I70A, P86K and L159A had reduced hydrolase activity in comparison to wild-type ATGL and responded to CGI-58 stimulation by partial increase of TGH activity. The third group with TGH activity comparable with wild-type included P103K, A104E, L178A, F348E and R351L. An intriguing observation was that the grouping of the ATGL mutants according to TGH activity correlated exactly with the type of switch mutants, *i.e.* the activity profiles traced to the impacted interaction partner. All three G0S2 switch mutants were inactive regardless of the presence of active CGI-58 (**Figure 4B** and **4C**, yellow label). The three Plin5 switch mutants, I70A, P86K and L159A, showed no basal activity and a moderate increase when stimulated with CGI-58 (**Figure 4BC**, dark blue label). The Plin1 switch mutants (**Figure 4BC** light blue label) and the CGI-58 switch mutant proteins (**Figure 4BC**, green label) were active, however the latter showed moderately reduced stimulated activity compared to wild-type ATGL. Therefore, we tested the CGI-58 switch mutants, F348E and R351L, for dose dependent simulation by increasing the concentration of affinity chromatography enriched human CGI-58 (**Figure 4D**). The experiment showed that basal TGH activity is not affected, however CGI-58 stimulation was significantly impaired under saturated conditions. In summary, we grouped ATGL switch variants according to defined patterns of *in vitro* TGH activities. This activity grouping strikingly resembled the selective interaction losses of individual partner proteins of the ATGL switch mutations.

### Lipolysis activity of ATGL switch variant in cells

We next used live cell confocal fluorescence microscopy to assay the activity of ATGL switch variants in mammalian cells. HeLa cells were transfected with YFP-tagged ATGL variants and then loaded with oleic acid for 6h, to induce LD formation. When stained with Bodipy, non-transfected cells showed a large number of LDs, while we readily observed distinct phenotypes of YFP positive cells expressing switch mutants (**Figure 5A**). Cells that expressed wild-type ATGL did not stain for LDs suggesting that ATGL overexpression had high lipolytic activity in live cells associated with strongly reduced TAG storage. In contrast a very high fraction of cells (>94%) expressing either of the three G0S2 switch ATGL variants contained LDs similar to non-transfected cells. For each switch mutant, we quantified the number of YFP positive cells for the presence of LDs reflecting lipolysis activity (**Figure 5B, Suppl. Table 6**). The results obtained with the ATGL variants agreed well with our THG activity data *in vitro*. The G0S2 switch variants were inactive *in vivo*, and also the PLIN5 switch variants I70A and P86K showed a high number of LD positive cells consistent with reduced *in vitro* activity. Conversely, ATGL L159A that showed reduced activity in the *in vitro* assay was active *in vivo*. The PLIN1 switch mutations showed full, wild-type like activity in cells. The CGI-58 switch variants, F348E and R351L were largely active under these conditions, again consistent with the TGH results. However quantitatively, a fraction of 6-9% LD containing cells indicated that the two switch variants with reduced binding of CGI-58 were functional, but exhibited reduced lipolytic activity. To test the hypothesis that the CGI-58 interaction site around amino acid position 350 in the C-terminal part of ATGL is critical for lipolysis we prepared an ATGL variant that combined both CGI-58 switch variants. The ATGL-F348E+R351L double mutant was inactive in the live cell lipolysis experiment (**Figure 5C**). Taken together, the live cell microscopy experiments revealed distinct cellular lipolysis phenotypes of ATGL variants which reflect the impact on the binding of ATGL to four interaction partners.

**Figure 5:**
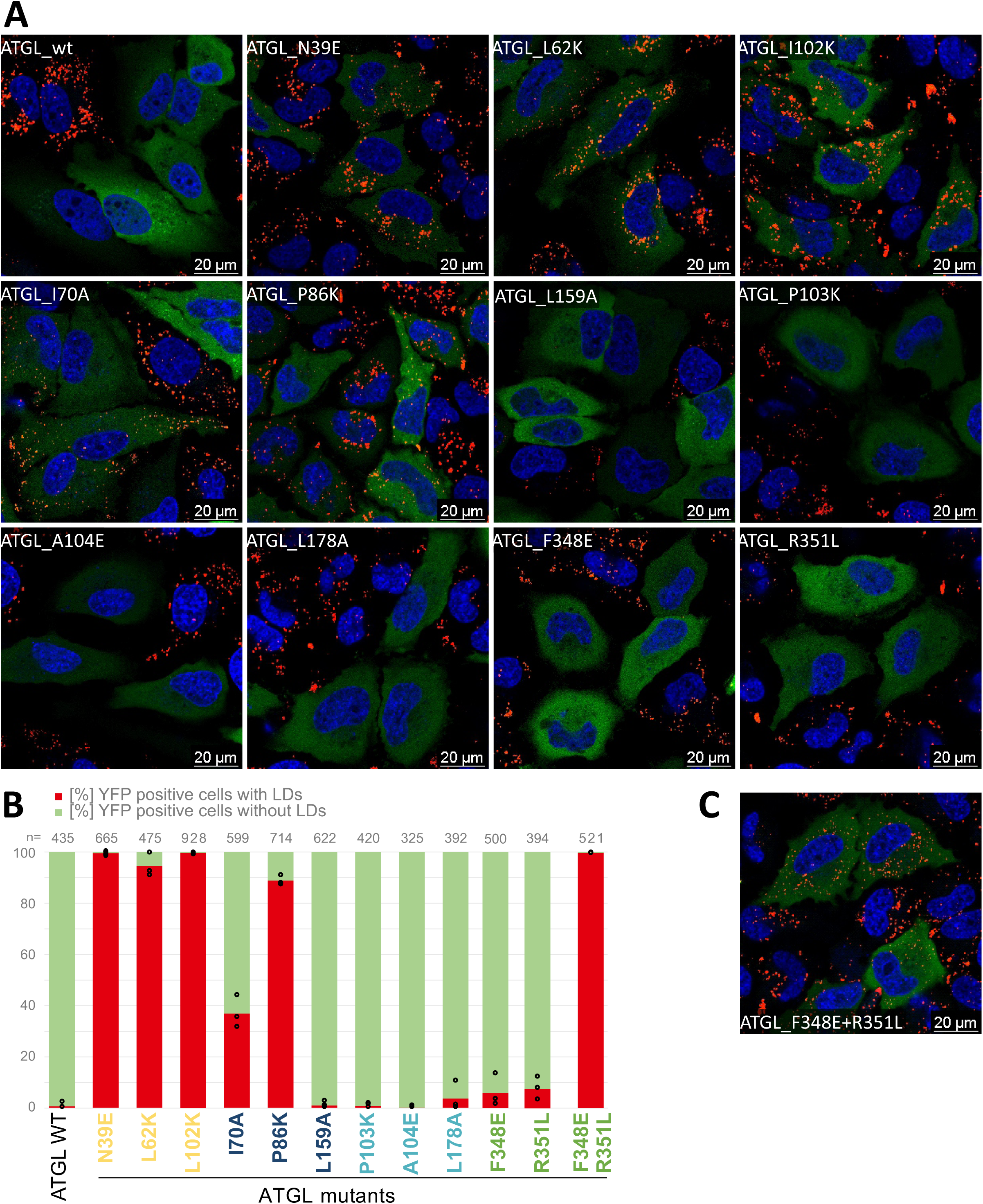
ATGL variant activity in live cells. **A.** Confocal images of HeLa cells transfected with YFP-ATGL variants. Pictures are merged fluorescent images with Hoechst in blue, YFP-ATGL in green and Bodipy in red. **B**. Fraction [%] of YFP positive cells with LDs (red) and without LDs (green). Cells expressing YFP-ATGL from three independent experiments were classified according to presence of absence (<2) of LDs. n = total number of cells counted. **C**. Confocal image of HeLa cells transfected with YFP-ATGL F348E+R351L double mutant. Picture as in A.

### ATGL variants inform about patient disease mutations

ATGL functional deficiency is associated with the rare neutral lipid storage disease with myopathy (NLSD-M) (Fischer et al., 2007). A few of the patients’ ATGL variants were functionally characterized. For example, E172K, P195L or R221P have reduced lipase activity while G483R does not properly localize to LDs (Tavian et al., 2012). ClinVar (Landrum et al., 2018) list 170 ATGL missense mutations found in patients with NLSD-M, all of which were classified as variants of unknown significance (VUS). Thirteen amino acid positions in the set of prioritized ATGL variants matched positions annotated in ClinVar. In contrast, we did not observe any overlap with population variants in ATGL (Karczewski et al., 2020) (**Figure 6A**). Importantly, the ClinVar overlap includes three of the switch mutants (P86K, A104E and R351L), a result that sheds a light on their function (**Figure 6A**). In our TGH assay, the P86K mutation was inactive under basal conditions, could only be stimulated partially and was inactive in cells, indicating that a substitution at this amino acid residue likely leads to an enzymatically impaired variant. While the amino acid substitutions A104E and R351L were enzymatically active, their interaction behavior is impaired, which likely results in altered ATGL function impacting NLSD. Taken together, these data demonstrate that single amino acid residues identified by deep mutational scanning can be useful to functionally annotate ATGL disease variants of unknown significance.

**Figure 6:**
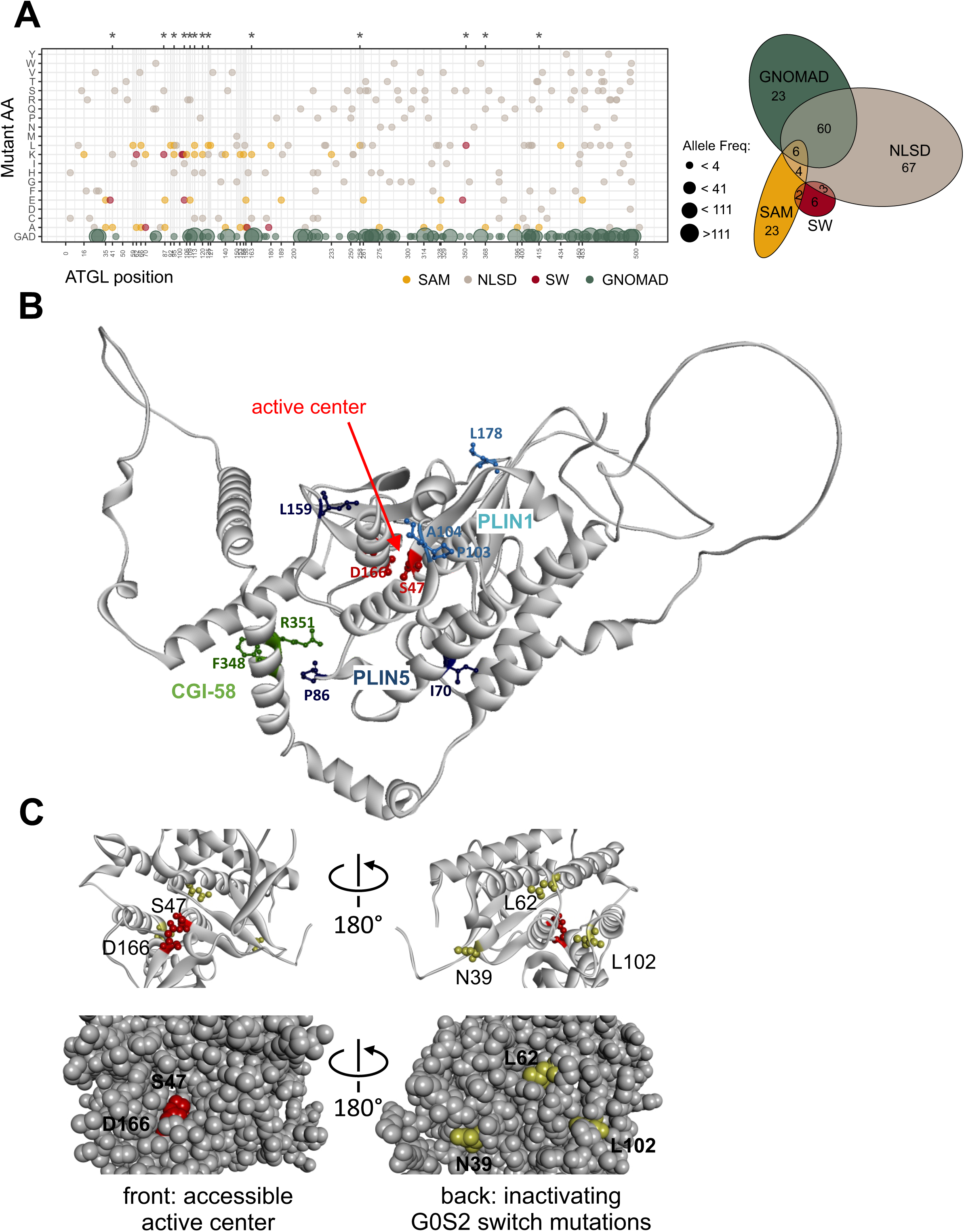
ATGL variants in a disease and structural context. **A.** Dotplot of annotated ATGL ClinVar missense substitutions associated with NLSD-M (ClinVar entries for PNPLA2, downloaded 10/2022). Gray dots: ClinVar mutations, yellow dots: selected ATGL mutations (SAM), red dots: switch mutations (SW). Green dots: variants from GnomAD (2.1.1. 03/2023). Asterisks on top highlight the positions T41, P86, R95, A104, H109, R113, R120, E125, R163, P258, R351, E368 and Y415, which overlap between the NLSD-M disease mutations and the SAM and SW mutants. *Right:* Euler diagram illustrating the overlaps of ClinVar missense mutations found in NLSD-M patients (gray), and the selected ATGL mutations (SAM, yellow) and the switch mutations (SW, red) and the missense variants from GnomAD (GNOMAD, green). **B.** AlphaFold 3D structure model of the ATGL C-backbone indicating binding partner contacts. Position and amino acid side chains (ball and stick) of *i)* the Plin5 switch variants I70, P86 and L159 (blue) and *ii)* of the Plin1 switch variants P103, A104 and L178 (cyan) and *iii)* of the CGI-58 switch variants F348 and R351 are displayed. The catalytic dyad comprising S44 and D166 (red) are highlighting the active center within the patatin domain fold. **C.** Zoom-in of the patatin domain. *Left:* view through the hydrophobic cavity towards the catalytic dyad comprising S44 and D166 (red) and the G0S2 switch positions N39, L62 and L102 (yellow) in the back. *Right:* structure turned by 180° around a vertical axis providing a view from the back of the domain, the side where the G0S2 switch mutations are solvent exposed. Bottom left, space fill model showing open cavity to the catalytic center. Bottom right, 180° turned view as above with the three switch mutations exposed to the surface in the back of the domain. The active center is not accessible from the side of the G0S2 switch residues. G0S2 switch positions L62 and L102 are located closest (∼15Å) to the catalytic residue S47.

### ATGL variants inform lipolysis mechanism

ATGL switch mutants that prevent the binding of a specific binding partner were not necessarily close in primary sequence. For example, the G0S2 switch variants: N39E, L62K and L102K span 60 amino acids in primary sequence. Likewise, PLIN1 switch mutations, P103K, A104E and L178A are more than 70 amino acids apart in primary sequence. Therefore, we projected the switch mutants on to available structural 3D models of ATGL (**Figure 6B**). Three-dimensional structure predictions of ATGL position the catalytic residues S47 and D166 in the center of the N-terminal patatin domain (Kulminskaya and Oberer, 2020) and additionally provide a low confident model for the unstructured C-terminal part (Jumper et al., 2021). The three Plin1 switch positions are spatially clustered together pointing at a distinct interaction site, while the Plin5 switch positions I70 and P86 localize on a different side of the patatin fold (**Figure 6B**). The Plin1 binding residues P103, A104 and L178 are surface residues with no intramolecular interactions of the side chains, in agreement with wild-type lipase activity of these ATGL variants. The side chains of Plin5 binding residues I70 and P86 were found to participate in intramolecular interactions (defined through a distance smaller than 5Å to another amino acid), providing a rational for our results as these mutations do both, prevent the PLIN5 interaction and affect the catalytic activity of ATGL.

Different ATGL binding determinants of the two perilipins are not unexpected. Plin5 is the major perilipin in the heart while Plin1 is highly expressed in adipose tissue, and knock-out as well as overexpression studies reveal different phenotypes (Saha et al., 2004; Kuramoto et al., 2012; Pollak et al., 2013; Itabe et al., 2017; Najt et al., 2022). Plin5 binds ATGL and CGI-58 via residues 200-463 in a mutually exclusive manner (Granneman et al., 2011). However, no ATGL residues important for perilipin binding are known. The binding mode of the ATGL-Plin1 interaction is elusive (Najt et al., 2022). Using chimeric and mutant Plin5 and Plin1 protein constructs, the perilipins were shown to bind ATGL differentially. In contrast to Plin5, IF and FRET experiments in COS7 cells could not demonstrate colocalization or interaction of Plin1 and ATGL at LDs (Granneman et al., 2011). In our analysis, co-IP experiments in HEK293T cells established a stable interaction of Plin1 and ATGL, that is characterized through three hydrolytically active Plin1 switch variants.

No ATGL binding determinants for the CGI-58 interaction are known either. We found variants affecting the interaction of ATGL with CGI-58 in the patatin domain as well as in the C-terminal part. Two CGI-58 switch mutations F348E and R351L were located in a structurally ill-defined C-terminal part of ATGL (**Figure 6B**). In dose dependent TGH activity measurements (**Fig. 4D**), the two switch variants showed reduced activity when stimulated with CGI-58. In live cells, we also observed an increased fraction of LD containing cells in comparison to wild-type ATGL (6-9%). Moreover, combining the two CGI-58 switch mutations resulted in an inactive ATGL variant with no lipolytic activity in cells. Therefore, we defined two highly specific CGI-58 switch variants outside of the patatin domain in the C-terminal part of ATGL previously not implicated in CGI-58 interaction and ATGL function.

It is well established that the N-terminal patatin domain extending to residue 254 of ATGL is sufficient for CGI-58 dependent stimulation of the lipid hydrolase activity (Schweiger et al., 2008; Cornaciu et al., 2011; Grabner et al., 2022). The C-terminal part of ATGL harbors the LD targeting sequence and phosphorylation sites including pT372, which abolishes LD localization (Xie et al., 2014) and pS406 which increases TAG hydrolase activity (Ahmadian et al., 2011). Schweiger et al. (Schweiger et al., 2008) compared TGH activity of full length ATGL protein with the truncated ATGL variant lacking 220 C-terminal amino acid residues. The truncated protein exhibited an up to 30-fold increased TGH activity upon CGI-58 mediated stimulation *in vitro*. Moreover, ATGL truncation increased binding to CGI-58 compared to full length ATGL by an unknown mechanism in a GST pulldown ELISA assay, leading the conclusion that the C-terminal part is a critical regulatory site for CGI-58 function (Schweiger et al., 2008). Like some missense mutations associated to ATGL malfunctioning, gene mutations resulting in premature stop codons also cause NLSD-M (NLSD1). ATGL patient mutations include truncations of C-terminal parts of ATGL, which leave the catalytically active patatin domain intact: Exon 5 I212X, Exon 5 L255X, Exon 7 L318X, Exon 7 L319X, Exon 7 Q289X (Schweiger et al., 2009). *In vivo* ATGL activity strongly depends on CGI-58 function as for example patient mutations in the CGI-58 gene (*ABHD5*) that lead to a decrease in ATGL activity, result in TAG accumulation causing NLSD-I (NLSD 2) (Schweiger et al., 2009). Overall, the functional and genetic data suggest that the 220 amino acid C-terminal part of ATGL has a crucial role for CGI-58 regulation of lipolysis activity. Here we defined switch residues in the region around amino acid position 350 that perturb the ATGL-CGI-58 interaction and are critical for cellular lipolysis. However, a full understanding of the role of the CGI-58 interaction with the C-terminus and the impact of the ATGL variants F348E and R351L in lipolysis requires further studies.

G0S2 is highly expressed in adipose tissue and differentiated adipocytes and constitutes the most potent inhibitor of ATGL activity (Schweiger et al., 2012; Zhang et al., 2017). G0S2 inhibits ATGL in a non-competitive manner (Cerk et al., 2014) and functions independently of CGI-58 (Yang et al., 2010; Cornaciu et al., 2011), leaving the binding mode of G0S2 with ATGL undefined. We identified a series of ATGL variants that strongly reduce or abolish G0S2 binding, including three G0S2 specific switch variants: N39E, L62K and L102K. Unexpectedly, all three were inactive in the TAG hydrolase activity assay, a finding that is strongly corroborated in our *in vivo* analysis. On the ATGL 3D structure model, these three residues were clustered together in proximity to the S44 and D166 catalytic dyad. The ATGL catalytic center is accessible through a cavity on the front side of the domain (**Figure 6C**). All three G0S2 switch mutations however clustered on the surface of the back side of the domain from where the active center is non-accessible. (**Figure 6C**, space fill model). This finding does not support a simple model where binding of G0S2 at or close to the substrate binding site directly perturbs substrate access or binding to the catalytic site. Rather it suggests that the GS02 switch mutations allosterically modulate the catalytic activity and that therefore G0S2 also inhibits the enzymatic activity of ATGL allosterically through binding at the back side of the patatin domain.

As GS02 switch mutations and G0S2 binding to ATGL strongly inhibit hydrolase activity, we suggest that the amino acid exchanges and the inhibitor binding act allosterically, both impairing conformational flexibility of the enzyme. The findings are strongly reminiscent of the (auto)inhibition mechanism of protein-tyrosine kinases (Hubbard et al., 1998). Protein kinases require conformational flexibility for phosphotransferase activity. Reduction of the conformational flexibility (both in its active or inactive conformation) through protein or small molecule binding inhibits kinase activity. For a large number of kinases this can be achieved through binding events at the back side of the kinase domain (e.g., SRC, ABL1), while the active site remains open and accessible (Huse and Kuriyan, 2002; Taipale et al., 2013). Recently, a reversible and competitive small molecule inhibitor (NG-497) for human ATGL was described which bound in the cavity near the active site involving residues 60–146 (Grabner et al., 2022). In agreement with our hypothesis that G0S2 binds the patatin domain on the back side, acting allosterically, the ATGL small molecule inhibitor and G0S2 were found to bind synergistically (Grabner et al., 2022).

## Conclusion

As the central protein with catalytic activity in lipolysis, ATGL interacts with a set of structurally diverse proteins. However, there is a lack of knowledge of ATGL binding determinants for its key regulatory interaction partners. To characterize the protein interactions with ATGL we applied comprehensive deep mutational interaction perturbation screening with five partner proteins. Mutations that affect binding to one protein typically have pleiotropic effects on binding to others (Sahni et al., 2015; Weng et al., 2022). However, here we defined a set of 11 switch mutations which are key in mediating interactions specifically to one of the partners with negligible effects on the others. Our results reveal a number of insights concerning the regulation of ATGL function. First and foremost, we defined distinct binding determinants for G0S2, CGI-58, Plin1 and Plin5 at the amino acid level, for which no such determinants have been reported previously. The switch mutations can be useful experimental tools, providing most selective perturbations to test the functions of individual interaction partners in various biological contexts (Woodsmith and Stelzl, 2017; Yadav et al., 2020). Second, we provide functional annotations for a set of 13 patient mutations associated to neutral lipid storage disease shedding light on their role in disease. Third, in agreement with their different functional roles in adipose and heart tissues, we identify distinct binding determinants for Plin1 and Plin5. Forth, our results provide strong hints for the importance of the ATGL C-terminal part for in the interaction with its activator CGI-58 and thus for cellular lipolysis. Fifth, the G0S2 switch variants of ATGL provide insight on the inhibition mechanism. The variants define an interaction site on the back of the patatin domain opposite of the catalytic center. The variants are no longer active suggesting an allosteric mode of action for G0S2 inhibition. To our knowledge, this deep mutational scanning study provides for the first-time amino acid resolution interaction surface information for ATGL’s partners.

## Material and Methods

### Clones

ORFs were obtained in the Gateway Entry vector or were amplified from a cDNA clone via PCR and inserted to an entry vector via BP reaction. The LR reaction was used to clone ORFs in Y2H vectors (Prey: pACT4-DM and pCBDU-JW; Bait: pBTM116-DM and pBTMcC24-DM) and for LUMIER assay in protein-A (pcDNA3.1PA-D57) and firefly- (pcDNA3.1V5-Fire) mammalian expression vectors (Worseck et al., 2012; Weimann et al., 2013). Reactions were performed using a standard protocol from Invitrogen. Swissprot Sequence IDs: ATGL(Q96AD5); mPLIN1(Q8CGN5); mPLIN5(Q8BVZ1); G0S2(P27469), CIDEC(Q96AQ7).

### Mutagenic library construction

On chip synthetized oligonucleotides were purchased from Cutsom Array, Inc. A total of 3692 ATGL primers were generated for each of the two libraries. The oligonucleotides (104 bp length: 60 bp primer + 44 bp adapter sequences) were amplified via PCR and adapter sequences were removed using restriction enzymes BciVI (New England BioLabs GmbH, R0596S) and BspQI (New England BioLabs GmbH, R0712S). Purified via an 8% acrylamide 8M urea gel. Bands of 60 bp size were cut out and gel pieces were incubated in 200 µl dH2O at 4°C for 24h to resolve the 60 bp oligonucleotides.

Deep mutagenesis protocol from E. Wrenbeck *et al*. was used (Wrenbeck et al., 2016). Minor changes were applied to increase the mutational power and reduce the WT-background (Plasmid Safe digest and DPNI digest).

### Interaction perturbation reverse Y2H screen

To ensure a highly efficient yeast transformation and thus a high coverage of all mutants in the DNA libraries the lithium acetate / single-stranded carrier DNA/ PEG method published by Gietz and Schießtl (Gietz and Schiestl, 2007) was used. For one ATGL library four aliquots, with 1.5 µg plasmid DNA each, were prepared. After transformation procedure the aliquots of the same DNA-libraries were united and 6ml NB media were added. 2.5ml of this suspension were plated on a large bioassay dish containing selective NB agar. The plates were incubated at 30°C for 4 days. The total colony number was calculated by counting a 1 cm^2^ area of grown colonies on 2 squares per plate.

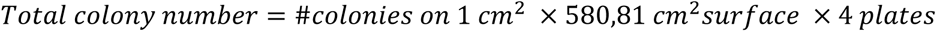

The yeast library transformants were used for the reverse Y2H-protocol which was published by Woodsmith *et al*. (Woodsmith et al., 2017) and performed accordingly. After the growth selection of the mated yeast strains, the grown colonies were collected, lysed and the DNA was purified through isopropanol precipitation and phenol-chloroform extraction and amplified via PCR.

### Data analysis

Next generation sequencing (Illumina NextSeq 500(SN442), NextSeq High PE151) including barcoding of PCR samples was performed at the Sequencing Core Facility of the Max-Planck-Institute (Berlin, Germany). The obtained sequencing data were processed as described previously (Woodsmith et al., 2017), a workflow incorporating Perl, as well as R scripts to analyze the paired-end sequencing data. In brief, fastq files of replicates were combined and converted to fasta files. The 150-mer reads were mapped to the wild-type ATGL cDNA sequence using the STAR Aligner. The amino acid codon enrichment was determined for each of the 18 protein interaction pairs tested.

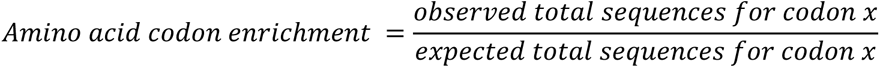

The cutoff for the number of reads with mutations, as well as the fold-linear model enrichment cutoff were chosen in a library specific manner (Suppl. Table 2). For non-coded mutations the cutoff for the number of reads was 10x the median codon sequences across all positions. In a final step a custom perl script was used to detect and remove secondary mutations.

### Site directed Mutagenesis

The site directed mutagenesis (Liu and Naismith, 2008) was performed using the Phusion High Fidelity DNA Polymerase (New England BioLabs GmbH, E0553L). 3.5ng template DNA as well as 5µM of each primer were used. PCR was performed for 30 cycles with an annealing temperature of 65.5°C. Subsequently the PCR samples (10µl) were digested with DPNI (0.2µl New England BioLabs GmbH, R0176S) in CutSmart Buffer to remove the wild-type background. The reaction was inactivated at 80°C for 20 minutes and competent TOP10 cells were transformed and utilized for plasmid preparation. Individual point mutations were verified via tag-sequencing.

### Luciferase based co-immunoprecipitation experiments in mammalian cells

IgG-coated LUMITRAC 96 well plates: Wells were incubated with sheep gamma globulin (Jackson ImmunoReasearch, 013-000-002) for 24 h at 4°C, followed by blocking with BSA () for another 24 h at 4°C. Finally, plates were incubated with AffiniPure Rabbit Anti-SheepIgG (H+L) (Jackson ImmunoResearch, 313-005-003) for 24 h at 4°C.

HEK293T cells were maintained in Dulbecco’s Modified Eagle’s Medium (DMEM) supplemented with 10% FBS and 1% Pen Strep at 37°C, 37.5% CO_2_ and 95% humidity. 2*10^4 HEK293T cells in a well of a 96-well plate (coated with 0.05 mg/ml poly-D-lysin, Sigma Aldrich, P7405) were transiently transfected with 80 ng of a pcDNA3.1PA-D57 construct and 80 ng of a pFireV5-DM construct using PEI (1mg/ml; Alfa Aesar, 900-98-6) as transfection reagent (ratio DNA:PEI = 1:5). After 48h cell lysis was performed with SDS-free RIPA2 buffer (50mM Tris HCl, 150mM NaCl, 1M EGTA, 1% NP40, 0.25% sodium deoxycholate, 50µg/ml dH_2_O and 1 cOmplete^TM^ mini-tablet per 10ml buffer) for 45 minutes at 4°C. Protein complexes precipitated from 80µl cleared cell-extract in IgG-coated LUMITRAC plates for 2h at 4°C. Plates were washed tree times with 100µl ice cold PBS. 40 µl PBS and 40µl Bright-Glo^TM^ luciferase assay reagent (Promega, E2650) were added per well. The luminescent signal of firefly luciferase activity was measured after ∼10 min at RT in a micro-plate reader (Counter Beckmann DTX800). The assay was performed in triplicates, the mean log2 raw luminescence signals and standard deviation for each plasmid DNA pair was determined. Within every microtiter plate and for each tested luciferase-fusion construct log2 values were normalized to background pA-fusion protein or wildtype pA-fusion protein binding, respectively. Ratios larger than two over background or smaller than 0.5 when compared to wild-type ATGL and a z-score larger than two were considered. Protein expression was validated by western blotting.

### *in vitro* TG hydrolase assay

The ATGL variants were expressed in Expi293 cells. 2.5×

10^6^ cells were transfected with 10.8µg DNA using the ExpiFectamine^TM^ 293 Transfection Kit (Thermo Fisher Scientific, A14525) according to manufacturer’s instructions. Cells were harvested after 48 hours, washed with PBS, resuspended in 4 ml HSL buffer (0.25M sucrose, 1mM EDTA, 1mM DTT, 1µg/ml pepstatin, 2µg/ml antipain and 20µg/ml leupeptin) and lysed by ultra-sonification (20% amplitude, 30 seconds). After centrifugation (14,000rpm, 4°C for 10 minutes), the soluble extract was adjusted to a total protein level of 3.5µg/µl. Protein expression was validated by western blotting. The *in vitro* radiolabeled triglyceride hydrolase activity assay was performed according to Schweiger *et al*. (Schweiger et al., 2014).

### Live cell confocal fluorescence microscopy

HeLa cells were maintained in high-glucose +/+ DMEM (Dulbecco’s Modified Eagle’s Medium supplemented with 1% PEN-STREP [Penicillin-Streptomycin] and 10% FBS [fetal bovine serum]) and incubated at 37°C, 37.5% CO2 and 95% humidity in a 24-well format (µ-Plate 24 Well Black ID 14mm ibiTreat, ibidi, 221205/5). HeLa cells were transfected with YFP-ATGL variants using Polyethylenimine (PEI, linear, M. W. 20,000 [Alfa Aesar, 9002-98-6]) as transfection reagent (ratio DNA:PEI = 1:5). After 18h incubation, cell were treated with 200µM oleic acid (Na-Oleat:BSA = 3.25:1 in PBS) for 6-8h. Cells were then stained with Hoechst 33342 (1:1000) and with Bodipy 493/503 (Thermo Fisher Scientific Inc., 2256834; f.c. 0.2µg/ml) for 30 min.

HeLa cells were imaged on a Stellaris 5 confocal microscope (Leica) using an HC PL APO 63x, 1.4 NA objective. Excitation was for Hoechst at 405nm, for Bodipy at 500 nm, and for YFP-ATGL at 516 nm. Emission was detected between 420 nm and 505 nm for Hoechst, 505 nm - 522 nm for Bodipy, and 545 nm - 625 nm for YFP-ATGL. Images were analyzed with LASX software package (Leica).

### ClinVar Data Analysis

All annotated ClinVar entries for ATGL were downloaded (ClinVar entries for PNPLA2, downloaded 10/2022). The entries were filtered for single amino acid substitutions leading to a total of 172 annotated SNVs, all associated with NLSD-M.

### Graphics

All basic graphic illustrations, if not stated otherwise, were generated using R, RStudio and the ggplot2 package.

## Data accessibility

Raw data are accessible via ENA Project Accession Number PRJEB60025. https://www.ebi.ac.uk/ena/browser/home.

## Supplemental Tables

provided as xlsx files.

Suppl. Table 1: Protein pairs in interaction perturbation screening

Suppl. Table 2: Cutoff values used in sequence data processing

Suppl. Table 3: Combined profiles (Figure 2A)

Suppl. Table 4: 52 ATGL single amino acid mutants

Suppl. Table 5: Co-IP results of the 52 ATGL single amino acid mutants (log2 fold change values, Figure 3)

Suppl. Table 6: Quantification of cells with LDs (Figure 5B)

## Acknowledgements

We thank Sandra Fasching and Sarah Masser for help with the experiments, Jonathan Woodsmith for support with data analyses and Natalia Kunowska for critical reading of the manuscript. We thank Bernd Timmermann and the members of MPI-MG Sequencing Facility (Berlin) for performing the second generation sequencing experiments. The work was funded by the Austrian Science Fund doc.fund Molecular Metabolism (DOC 50). The work was also supported by the FWF (project P30162) and the Field of Excellence BioHealth-University of Graz.

## Author contributions

Conceptualization: US

Data curation: JMK, BH

Formal Analysis: JMK, BH

Funding acquisition: US, RZ

Investigation: JMK, GFG, AN, AH, EJ-L

Methodology: JMK, GFG, HE, RZ, US

Project administration:

Resources: US, RZ

Software:

Supervision: US, RZ, HE

Validation: JMK, GFG

Visualization: JMK, GFG, BH, US

Writing – original draft: US, JMK

Writing – review & editing: US, JMK and all authors

## References

Ahmadian, M., Abbott, M.J., Tang, T., Hudak, C.S.S., Kim, Y., and Bruss, M., et al. (2011). Desnutrin/ATGL is regulated by AMPK and is required for a brown adipose phenotype. Cell metabolism 13, 739–748, [10.1016/j.cmet.2011.05.002].

Cerk, I.K., Salzburger, B., Boeszoermenyi, A., Heier, C., Pillip, C., and Romauch, M., et al. (2014). A peptide derived from G0/G1 switch gene 2 acts as noncompetitive inhibitor of adipose triglyceride lipase. The Journal of biological chemistry 289, 32559–32570, [10.1074/jbc.M114.602599].

Cerk, I.K., Wechselberger, L., and Oberer, M. (2018). Adipose Triglyceride Lipase Regulation: An Overview. Current protein & peptide science 19, 221–233, [10.2174/1389203718666170918160110].

Cornaciu, I., Boeszoermenyi, A., Lindermuth, H., Nagy, H.M., Cerk, I.K., and Ebner, C., et al. (2011). The minimal domain of adipose triglyceride lipase (ATGL) ranges until leucine 254 and can be activated and inhibited by CGI-58 and G0S2, respectively. PloS one 6, e26349, [10.1371/journal.pone.0026349].

Diss, G., and Lehner, B. (2018). The genetic landscape of a physical interaction. eLife 7, [10.7554/eLife.32472].

Faure, A.J., Domingo, J., Schmiedel, J.M., Hidalgo-Carcedo, C., Diss, G., and Lehner, B. (2022). Mapping the energetic and allosteric landscapes of protein binding domains. Nature 604, 175–183, [10.1038/s41586-022-04586-4].

Fischer, J., Lefèvre, C., Morava, E., Mussini, J.-M., Laforêt, P., and Negre-Salvayre, A., et al. (2007). The gene encoding adipose triglyceride lipase (PNPLA2) is mutated in neutral lipid storage disease with myopathy. Nature genetics 39, 28–30, [10.1038/ng1951].

Gietz, R.D., and Schiestl, R.H. (2007). High-efficiency yeast transformation using the LiAc/SS carrier DNA/PEG method. Nature protocols 2, 31–34, [10.1038/nprot.2007.13].

Grabner, G.F., Guttenberger, N., Mayer, N., Migglautsch-Sulzer, A.K., Lembacher-Fadum, C., and Fawzy, N., et al. (2022). Small-Molecule Inhibitors Targeting Lipolysis in Human Adipocytes. Journal of the American Chemical Society 144, 6237–6250, [10.1021/jacs.1c10836].

Grabner, G.F., Xie, H., Schweiger, M., and Zechner, R. (2021). Lipolysis: cellular mechanisms for lipid mobilization from fat stores. Nature metabolism 3, 1445–1465, [10.1038/s42255-021-00493-6].

Grahn, T.H.M., Kaur, R., Yin, J., Schweiger, M., Sharma, V.M., and Lee, M.-J., et al. (2014). Fat-specific protein 27 (FSP27) interacts with adipose triglyceride lipase (ATGL) to regulate lipolysis and insulin sensitivity in human adipocytes. The Journal of biological chemistry 289, 12029–12039, [10.1074/jbc.M113.539890].

Granneman, J.G., Moore, H.-P.H., Krishnamoorthy, R., and Rathod, M. (2009). Perilipin controls lipolysis by regulating the interactions of AB-hydrolase containing 5 (Abhd5) and adipose triglyceride lipase (Atgl). The Journal of biological chemistry 284, 34538–34544, [10.1074/jbc.M109.068478].

Granneman, J.G., Moore, H.-P.H., Mottillo, E.P., Zhu, Z., and Zhou, L. (2011). Interactions of perilipin-5 (Plin5) with adipose triglyceride lipase. The Journal of biological chemistry 286, 5126–5135, [10.1074/jbc.M110.180711].

Haemmerle, G., Lass, A., Zimmermann, R., Gorkiewicz, G., Meyer, C., and Rozman, J., et al. (2006). Defective lipolysis and altered energy metabolism in mice lacking adipose triglyceride lipase. Science (New York, N.Y.) 312, 734–737, [10.1126/science.1123965].

Haemmerle, G., Moustafa, T., Woelkart, G., Büttner, S., Schmidt, A., and van de Weijer, T., et al. (2011). ATGL-mediated fat catabolism regulates cardiac mitochondrial function via PPAR-α and PGC-1. Nature medicine 17, 1076–1085, [10.1038/nm.2439].

Hegele, A., Kamburov, A., Grossmann, A., Sourlis, C., Wowro, S., and Weimann, M., et al. (2012). Dynamic protein-protein interaction wiring of the human spliceosome. Molecular cell 45, 567–580, [10.1016/j.molcel.2011.12.034.].

Hidalgo, F., Nocka, L.M., Shah, N.H., Gorday, K., Latorraca, N.R., and Bandaru, P., et al. (2022). A saturation-mutagenesis analysis of the interplay between stability and activation in Ras. eLife 11, [10.7554/eLife.76595].

Hubbard, S.R., Mohammadi, M., and Schlessinger, J. (1998). Autoregulatory mechanisms in protein-tyrosine kinases. Journal of Biological Chemistry 273, 11987–11990, [10.1074/jbc.273.20.11987].

Huse, M., and Kuriyan, J. (2002). The conformational plasticity of protein kinases. Cell 109, 275–282, [10.1016/S0092-8674(02)00741-9].

Itabe, H., Yamaguchi, T., Nimura, S., and Sasabe, N. (2017). Perilipins: a diversity of intracellular lipid droplet proteins. Lipids in health and disease 16, 83, [10.1186/s12944-017-0473-y].

Jumper, J., Evans, R., Pritzel, A., Green, T., Figurnov, M., and Ronneberger, O., et al. (2021). Highly accurate protein structure prediction with AlphaFold. Nature 596, 583–589, [10.1038/s41586-021-03819-2].

Karczewski, K.J., Francioli, L.C., Tiao, G., Cummings, B.B., Alföldi, J., and Wang, Q., et al. (2020). The mutational constraint spectrum quantified from variation in 141,456 humans. Nature 581, 434–443, [10.1038/s41586-020-2308-7].

Kien, B., Kolleritsch, S., Kunowska, N., Heier, C., Chalhoub, G., and Tilp, A., et al. (2022). Lipid droplet-mitochondria coupling via perilipin 5 augments respiratory capacity but is dispensable for FA oxidation. Journal of lipid research 63, 100172, [10.1016/j.jlr.2022.100172].

Kulminskaya, N., and Oberer, M. (2020). Protein-protein interactions regulate the activity of Adipose Triglyceride Lipase in intracellular lipolysis. Biochimie 169, 62–68, [10.1016/j.biochi.2019.08.004].

Kunowska, N., and Stelzl, U. (2022). Decoding the cellular effects of genetic variation through interaction proteomics. Current opinion in chemical biology 66, 102100, [10.1016/j.cbpa.2021.102100].

Kuramoto, K., Okamura, T., Yamaguchi, T., Nakamura, T.Y., Wakabayashi, S., and Morinaga, H., et al. (2012). Perilipin 5, a lipid droplet-binding protein, protects heart from oxidative burden by sequestering fatty acid from excessive oxidation. The Journal of biological chemistry 287, 23852–23863, [10.1074/jbc.M111.328708].

Landrum, M.J., Lee, J.M., Benson, M., Brown, G.R., Chao, C., and Chitipiralla, S., et al. (2018). ClinVar: improving access to variant interpretations and supporting evidence. Nucleic acids research 46, D1062–D1067, [10.1093/nar/gkx1153].

Lass, A., Zimmermann, R., Haemmerle, G., Riederer, M., Schoiswohl, G., and Schweiger, M., et al. (2006). Adipose triglyceride lipase-mediated lipolysis of cellular fat stores is activated by CGI-58 and defective in Chanarin-Dorfman Syndrome. Cell metabolism 3, 309–319, [10.1016/j.cmet.2006.03.005].

Liu, H., and Naismith, J.H. (2008). An efficient one-step site-directed deletion, insertion, single and multiple-site plasmid mutagenesis protocol. BMC biotechnology 8, 91, [10.1186/1472-6750-8-91].

Miyoshi, H., Souza, S.C., Endo, M., Sawada, T., Perfield, J.W., and Shimizu, C., et al. (2010). Perilipin overexpression in mice protects against diet-induced obesity. Journal of lipid research 51, 975–982, [10.1194/jlr.M002352].

Moesslacher, C.S., Kohlmayr, J.M., and Stelzl, U. (2021). Exploring absent protein function in yeast: assaying post translational modification and human genetic variation. Microb Cell 8, 164–183, [10.15698/mic2021.08.756].

Moesslacher, C.S., Woodsmith, J., Auernig, E., Feichtner, A., Jany-Luig, E., and Jehle, S., et al. (2022). Missense variant interaction scanning reveals a critical role of the FERM-F3 domain for tumor suppressor protein NF2 conformation and function, [10.1101/2022.12.11.519953].

Najt, C.P., Devarajan, M., and Mashek, D.G. (2022). Perilipins at a glance. Journal of cell science 135, [10.1242/jcs.259501].

Olzmann, J.A., and Carvalho, P. (2019). Dynamics and functions of lipid droplets. Nature reviews. Molecular cell biology 20, 137–155, [10.1038/s41580-018-0085-z].

Pollak, N.M., Jaeger, D., Kolleritsch, S., Zimmermann, R., Zechner, R., and Lass, A., et al. (2015). The interplay of protein kinase A and perilipin 5 regulates cardiac lipolysis. The Journal of biological chemistry 290, 1295–1306, [10.1074/jbc.M114.604744].

Pollak, N.M., Schweiger, M., Jaeger, D., Kolb, D., Kumari, M., and Schreiber, R., et al. (2013). Cardiac-specific overexpression of perilipin 5 provokes severe cardiac steatosis via the formation of a lipolytic barrier. Journal of lipid research 54, 1092–1102, [10.1194/jlr.M034710].

Puri, V., Konda, S., Ranjit, S., Aouadi, M., Chawla, A., and Chouinard, M., et al. (2007). Fat-specific protein 27, a novel lipid droplet protein that enhances triglyceride storage. The Journal of biological chemistry 282, 34213–34218, [10.1074/jbc.M707404200].

Recazens, E., Mouisel, E., and Langin, D. (2021). Hormone-sensitive lipase: sixty years later. Progress in lipid research 82, 101084, [10.1016/j.plipres.2020.101084].

Riegler-Berket, L., Wechselberger, L., Cerk, I.K., Padmanabha Das, K.M., Viertlmayr, R., and Kulminskaya, N., et al. (2022). Residues of the minimal sequence of G0S2 collectively contribute to ATGL inhibition while C-and N-terminal extensions promote binding to ATGL. Biochimica et biophysica acta. Molecular and cell biology of lipids 1867, 159105, [10.1016/j.bbalip.2021.159105].

Rondini, E.A., Mladenovic-Lucas, L., Roush, W.R., Halvorsen, G.T., Green, A.E., and Granneman, J.G. (2017). Novel Pharmacological Probes Reveal ABHD5 as a Locus of Lipolysis Control in White and Brown Adipocytes. The Journal of pharmacology and experimental therapeutics 363, 367–376, [10.1124/jpet.117.243253].

Saha, P.K., Kojima, H., Martinez-Botas, J., Sunehag, A.L., and Chan, L. (2004). Metabolic adaptations in the absence of perilipin: increased beta-oxidation and decreased hepatic glucose production associated with peripheral insulin resistance but normal glucose tolerance in perilipin-null mice. Journal of Biological Chemistry 279, 35150–35158, [10.1074/jbc.M405499200].

Sahni, N., Yi, S., Taipale, M., Fuxman Bass, J.I., Coulombe-Huntington, J., and Yang, F., et al. (2015). Widespread macromolecular interaction perturbations in human genetic disorders. Cell 161, 647–660, [10.1016/j.cell.2015.04.013].

Sahu-Osen, A., Montero-Moran, G., Schittmayer, M., Fritz, K., Dinh, A., and Chang, Y.-F., et al. (2015). CGI-58/ABHD5 is phosphorylated on Ser239 by protein kinase A: control of subcellular localization. Journal of lipid research 56, 109–121, [10.1194/jlr.M055004].

Sawada, T., Miyoshi, H., Shimada, K., Suzuki, A., Okamatsu-Ogura, Y., and Perfield, J.W., et al. (2010). Perilipin overexpression in white adipose tissue induces a brown fat-like phenotype. PloS one 5, e14006, [10.1371/journal.pone.0014006].

Schoiswohl, G., Schweiger, M., Schreiber, R., Gorkiewicz, G., Preiss-Landl, K., and Taschler, U., et al. (2010). Adipose triglyceride lipase plays a key role in the supply of the working muscle with fatty acids. Journal of lipid research 51, 490–499, [10.1194/jlr.M001073].

Schoiswohl, G., Stefanovic-Racic, M., Menke, M.N., Wills, R.C., Surlow, B.A., and Basantani, M.K., et al. (2015). Impact of Reduced ATGL-Mediated Adipocyte Lipolysis on Obesity-Associated Insulin Resistance and Inflammation in Male Mice. Endocrinology 156, 3610–3624, [10.1210/en.2015-1322].

Schreiber, R., Hofer, P., Taschler, U., Voshol, P.J., Rechberger, G.N., and Kotzbeck, P., et al. (2015). Hypophagia and metabolic adaptations in mice with defective ATGL-mediated lipolysis cause resistance to HFD-induced obesity. Proceedings of the National Academy of Sciences of the United States of America 112, 13850–13855, [10.1073/pnas.1516004112].

Schweiger, M., Eichmann, T.O., Taschler, U., Zimmermann, R., Zechner, R., and Lass, A. (2014). Measurement of lipolysis. Methods in enzymology 538, 171–193, [10.1016/B978-0-12-800280-3.00010-4].

Schweiger, M., Lass, A., Zimmermann, R., Eichmann, T.O., and Zechner, R. (2009). Neutral lipid storage disease: genetic disorders caused by mutations in adipose triglyceride lipase/PNPLA2 or CGI-58/ABHD5. American journal of physiology. Endocrinology and metabolism 297, E289–96, [10.1152/ajpendo.00099.2009].

Schweiger, M., Paar, M., Eder, C., Brandis, J., Moser, E., and Gorkiewicz, G., et al. (2012). G0/G1 switch gene-2 regulates human adipocyte lipolysis by affecting activity and localization of adipose triglyceride lipase. Journal of lipid research 53, 2307–2317, [10.1194/jlr.M027409].

Schweiger, M., Romauch, M., Schreiber, R., Grabner, G.F., Hütter, S., and Kotzbeck, P., et al. (2017). Pharmacological inhibition of adipose triglyceride lipase corrects high-fat diet-induced insulin resistance and hepatosteatosis in mice. Nature communications 8, 14859, [10.1038/ncomms14859].

Schweiger, M., Schoiswohl, G., Lass, A., Radner, F.P.W., Haemmerle, G., and Malli, R., et al. (2008). The C-terminal region of human adipose triglyceride lipase affects enzyme activity and lipid droplet binding. The Journal of biological chemistry 283, 17211–17220, [10.1074/jbc.M710566200].

Shendure, J., and Fields, S. (2016). Massively Parallel Genetics. Genetics 203, 617–619, [10.1534/genetics.115.180562].

Starita, L.M., Young, D.L., Islam, M., Kitzman, J.O., Gullingsrud, J., and Hause, R.J., et al. (2015). Massively Parallel Functional Analysis of BRCA1 RING Domain Variants. Genetics 200, 413–422, [10.1534/genetics.115.175802].

Subramanian, V., Rothenberg, A., Gomez, C., Cohen, A.W., Garcia, A., and Bhattacharyya, S., et al. (2004). Perilipin A mediates the reversible binding of CGI-58 to lipid droplets in 3T3-L1 adipocytes. The Journal of biological chemistry 279, 42062–42071, [10.1074/jbc.M407462200].

Taipale, M., Krykbaeva, I., Whitesell, L., Santagata, S., Zhang, J., and Liu, Q., et al. (2013). Chaperones as thermodynamic sensors of drug-target interactions reveal kinase inhibitor specificities in living cells. Nature biotechnology 31, 630–637, [10.1038/nbt.2620].

Tansey, J.T., Sztalryd, C., Gruia-Gray, J., Roush, D.L., Zee, J.V., and Gavrilova, O., et al. (2001). Perilipin ablation results in a lean mouse with aberrant adipocyte lipolysis, enhanced leptin production, and resistance to diet-induced obesity. Proceedings of the National Academy of Sciences of the United States of America 98, 6494–6499, [10.1073/pnas.101042998].

Tavian, D., Missaglia, S., Redaelli, C., Pennisi, E.M., Invernici, G., and Wessalowski, R., et al. (2012). Contribution of novel ATGL missense mutations to the clinical phenotype of NLSD-M: a strikingly low amount of lipase activity may preserve cardiac function. Human molecular genetics 21, 5318–5328, [10.1093/hmg/dds388].

Wang, H., Sreenivasan, U., Hu, H., Saladino, A., Polster, B.M., and Lund, L.M., et al. (2011). Perilipin 5, a lipid droplet-associated protein, provides physical and metabolic linkage to mitochondria. Journal of lipid research 52, 2159–2168, [10.1194/jlr.M017939].

Weimann, M., Grossmann, A., Woodsmith, J., Özkan, Z., Birth, P., and Meierhofer, D., et al. (2013). A Y2H-seq approach defines the human protein methyltransferase interactome. Nat Methods 10, 339–342, [10.1038/nmeth.2397].

Weng, C., Faure, A.J., and Lehner, B. (2022). The energetic and allosteric landscape for KRAS inhibition, [10.1101/2022.12.06.519122].

Woodsmith, J., Apelt, L., Casado-Medrano, V., Özkan, Z., Timmermann, B., and Stelzl, U. (2017). Protein interaction perturbation profiling at amino-acid resolution. Nat Methods 14, 1213–1221, [10.1038/nmeth.4464].

Woodsmith, J., and Stelzl, U. (2017). Understanding Disease Variants through the Lens of Protein Interactions. Cell systems 5, 544–546, [10.1016/j.cels.2017.12.009].

Worseck, J.M., Grossmann, A., Weimann, M., Hegele, A., and Stelzl, U. (2012). A stringent yeast two-hybrid matrix screening approach for protein-protein interaction discovery. Methods in molecular biology (Clifton, N.J.) 812, 63–87, [10.1007/978-1-61779-455-1_4].

Wrenbeck, E.E., Klesmith, J.R., Stapleton, J.A., Adeniran, A., Tyo, K.E.J., and Whitehead, T.A. (2016). Plasmid-based one-pot saturation mutagenesis. Nat Methods 13, 928–930, [10.1038/nmeth.4029].

Xie, X., Langlais, P., Zhang, X., Heckmann, B.L., Saarinen, A.M., and Mandarino, L.J., et al. (2014). Identification of a novel phosphorylation site in adipose triglyceride lipase as a regulator of lipid droplet localization. American journal of physiology. Endocrinology and metabolism 306, E1449–59, [10.1152/ajpendo.00663.2013].

Yadav, A., Vidal, M., and Luck, K. (2020). Precision medicine - networks to the rescue. Current opinion in biotechnology 63, 177–189, [10.1016/j.copbio.2020.02.005].

Yamaguchi, T., Omatsu, N., Matsushita, S., and Osumi, T. (2004). CGI-58 interacts with perilipin and is localized to lipid droplets. Possible involvement of CGI-58 mislocalization in Chanarin-Dorfman syndrome. The Journal of biological chemistry 279, 30490–30497, [10.1074/jbc.M403920200].

Yang, X., Heckmann, B.L., Zhang, X., Smas, C.M., and Liu, J. (2013). Distinct mechanisms regulate ATGL-mediated adipocyte lipolysis by lipid droplet coat proteins. Molecular endocrinology (Baltimore, Md.) 27, 116–126, [10.1210/me.2012-1178].

Yang, X., Lu, X., Lombès, M., Rha, G.B., Chi, Y.-I., and Guerin, T.M., et al. (2010). The G(0)/G(1) switch gene 2 regulates adipose lipolysis through association with adipose triglyceride lipase. Cell metabolism 11, 194–205, [10.1016/j.cmet.2010.02.003].

Zhang, X., Heckmann, B.L., Campbell, L.E., and Liu, J. (2017). G0S2: A small giant controller of lipolysis and adipose-liver fatty acid flux. Biochimica et biophysica acta. Molecular and cell biology of lipids 1862, 1146–1154, [10.1016/j.bbalip.2017.06.007].

Zimmermann, R., Strauss, J.G., Haemmerle, G., Schoiswohl, G., Birner-Gruenberger, R., and Riederer, M., et al. (2004). Fat mobilization in adipose tissue is promoted by adipose triglyceride lipase. Science (New York, N.Y.) 306, 1383–1386, [10.1126/science.1100747].

